# SharePro: an accurate and efficient genetic colocalization method accounting for multiple causal signals

**DOI:** 10.1101/2023.07.24.550431

**Authors:** Wenmin Zhang, Tianyuan Lu, Robert Sladek, Yue Li, Hamed S. Najafabadi, Josée Dupuis

**Affiliations:** Quantitative Life Sciences, McGill University, Montreal, Canada; Department of Statistical Sciences, University of Toronto, Toronto, Canada; Department of Human Genetics, McGill University, Montreal, Canada; Dahdaleh Institute of Genomic Medicine, Montreal, Canada; School of Computer Science, McGill University, Montreal, Canada; Department of Epidemiology, Biostatistics and Occupational Health, McGill University, Montreal, Canada

## Abstract

**Motivation:** Colocalization analysis is commonly used to assess whether two or more traits share the same genetic signals identified in genome-wide association studies (GWAS), and is important for prioritizing targets for functional follow-up of GWAS results. Existing colocalization methods can have suboptimal performance when there are multiple causal variants in one genomic locus.

**Results:** We propose SharePro to extend the COLOC framework for colocalization analysis. SharePro integrates linkage disequilibrium (LD) modelling and colocalization assessment by grouping correlated variants into effect groups. With an efficient variational inference algorithm, posterior colocalization probabilities can be accurately estimated. In simulation studies, SharePro demonstrated increased power with a well-controlled false positive rate at a low computational cost. Through a challenging case of the colocalization analysis of the circulating abundance of R-spondin 3 (RSPO3) GWAS and estimated bone mineral density GWAS, we demonstrated the utility of SharePro in identifying biologically plausible colocalized signals.

**Availability and Implementation:** The SharePro software for colocalization analysis is openly available at https://github.com/zhwm/SharePro_coloc and the analysis conducted in this study is available at https://github.com/zhwm/SharePro_coloc_analysis.

## 1 Introduction

Colocalization analysis is a commonly used statistical procedure to assess whether two or more traits share the same genetic signals identified in genome-wide association studies (GWAS) [1–5]. It is important for understanding the interplay between heritable traits [6,7], such as validating causal inference results based on Mendelian randomization analysis [3, 8, 9] and identifying candidate genes for functional follow-up studies [2, 10–12]. Therefore, a sensitive colocalization method that effectively controls the false positive rate is crucial for increasing the yield of complex trait genetics studies.

COLOC [1] is one of the most widely used methods for colocalization analysis. COLOC uses a Bayesian framework to estimate posterior probabilities of five different causal settings in a locus (H0: no causal signals; H1: one unique causal signal for trait 1; H2: one unique causal signal for trait 2; H3: different causal signals for trait 1 and 2; H4: one shared causal signal for trait 1 and 2. [1]). Colocalization probability is defined by the posterior probability of H4 [1]. A key assumption in COLOC is that only one causal variant exists within each genomic locus [1]. In both simulation and substantive studies [1,10], COLOC demonstrated high accuracy in identifying the shared causal signal when the one-causal-variant assumption was met. However, the performance of COLOC may be compromised when more than one causal signal exists in a genomic locus [2, 5, 13].

Building upon COLOC, several methods have been developed to address these challenges. For example, eCAVIAR allows for multiple causal signals [2] by adopting the CAVIAR [14] fine-mapping framework for colocalization. In eCAVIAR, colocalization is assessed at the variant-level by examining the probabilities of variants being causal in both traits. Specifically, the posterior inclusion probabilities for variants are first calculated separately for each trait. Then, the variant-level colocalization probabilities are obtained from the product of the posterior inclusion probabilities. Recently, COLOC + SuSiE [5] adopts a fine-mapping method SuSiE [15] for identifying multiple causal variants before performing pairwise colocalization, which could improve the performance of COLOC when multiple causal signals exist. Similarly, PWCoCo [16] first performs conditional and joint analysis with GCTA-COJO [17], followed by colocalization analysis on each pair of the conditionally independent signals identified by GCTA-COJO using COLOC. These methods implement a two-step strategy. Namely, they first account for LD via fine-mapping or conditional analysis to identify candidate variants for colocalization analysis, separately for each trait. And then, under the one-causal-variant assumption, colocalization probabilities are assessed by examining whether each pair of candidate variants represents the same causal signal. However, with this strategy, the uncertainties in accounting for LD from the first step might affect the assessment of colocalization in the second step.

We propose SharePro (Shared sparse Projection for colocalization analysis) to integrate LD modelling and colocalization assessment to account for multiple causal variants in colocalization analysis. In SharePro, highly correlated variants are grouped into effect groups and colocalization probabilities are assessed by examining the causal status of each effect group in different traits. We evaluate the performance of SharePro in simulation studies in comparison to state-of-the-art colocalization methods. We further examine colocalization between cis-protein quantitative trait locus (pQTL) of the circulating abundance of RSPO3 and a GWAS locus identified for estimated bone mineral density (eBMD) using heel quantitative ultrasound measurement to evaluate whether SharePro could better identify biologically plausible colocalized signals.

## 2 Methods

### 2.1 SharePro method overview

SharePro takes marginal associations (z-scores) from GWAS summary statistics and LD information calculated from a reference panel as inputs, and infers posterior probabilities of colocalization (**Figure 1**). Unlike existing methods, SharePro takes an effect group-level approach for colocalization. Specifically, SharePro uses a sparse projection shared across traits to group correlated variants into effect groups. Through this shared projection, variant representations for effect groups are the same across traits so that colocalization probabilities can be directly calculated at the effect group-level. With an efficient variational inference algorithm, both variant representations for effect groups and their causal statuses in traits can be accurately inferred. Consequently, we can obtain colocalization probabilities from the posterior probabilities of effect groups being causal for all traits.

**Figure 1.**
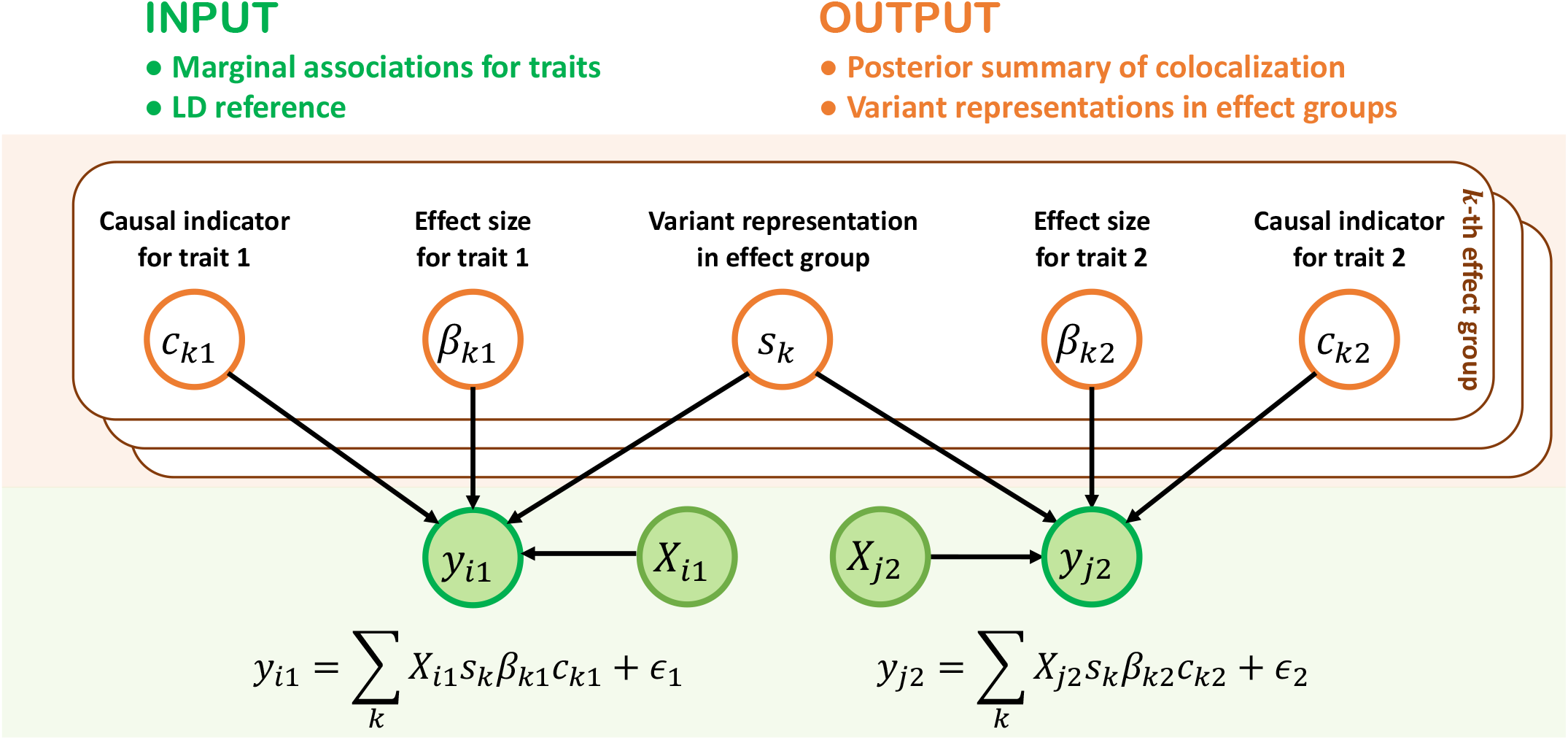
SharePro for genetic colocalization analysis. The data generative process in SharePro is depicted in the graphical model. Green shaded nodes represent observed variables: genotype *X*_*i*1_, trait *y*_*i*1_ for the *i*^*th*^ individual in the first study, and genotype *X*_*j*2_, trait *y*_*j*2_ for the *j*^*th*^ individual in the second study. The orange unshaded nodes represent latent variables characterizing effect groups. **s**_*k*_ is a sparse indicator shared between traits, specifying variant representations for the *k*^*th*^ effect group. *c*_*k*1_ and *c*_*k*2_ are causal indicators of whether the *k*^*th*^ effect group is causal in trait **y**_1_ and trait **y**_2_ while *β*_*k*1_ and *β*_*k*2_ represent the corresponding effect sizes. As a result, colocalization probability for the *k*^*th*^ effect group is the posterior probability of *c*_*k*1_ = *c*_*k*2_ = 1. Here we assume individual-level data are available and adaption to GWAS summary statistics is detailed in the **Supplementary Notes**.

**Figure 2.**
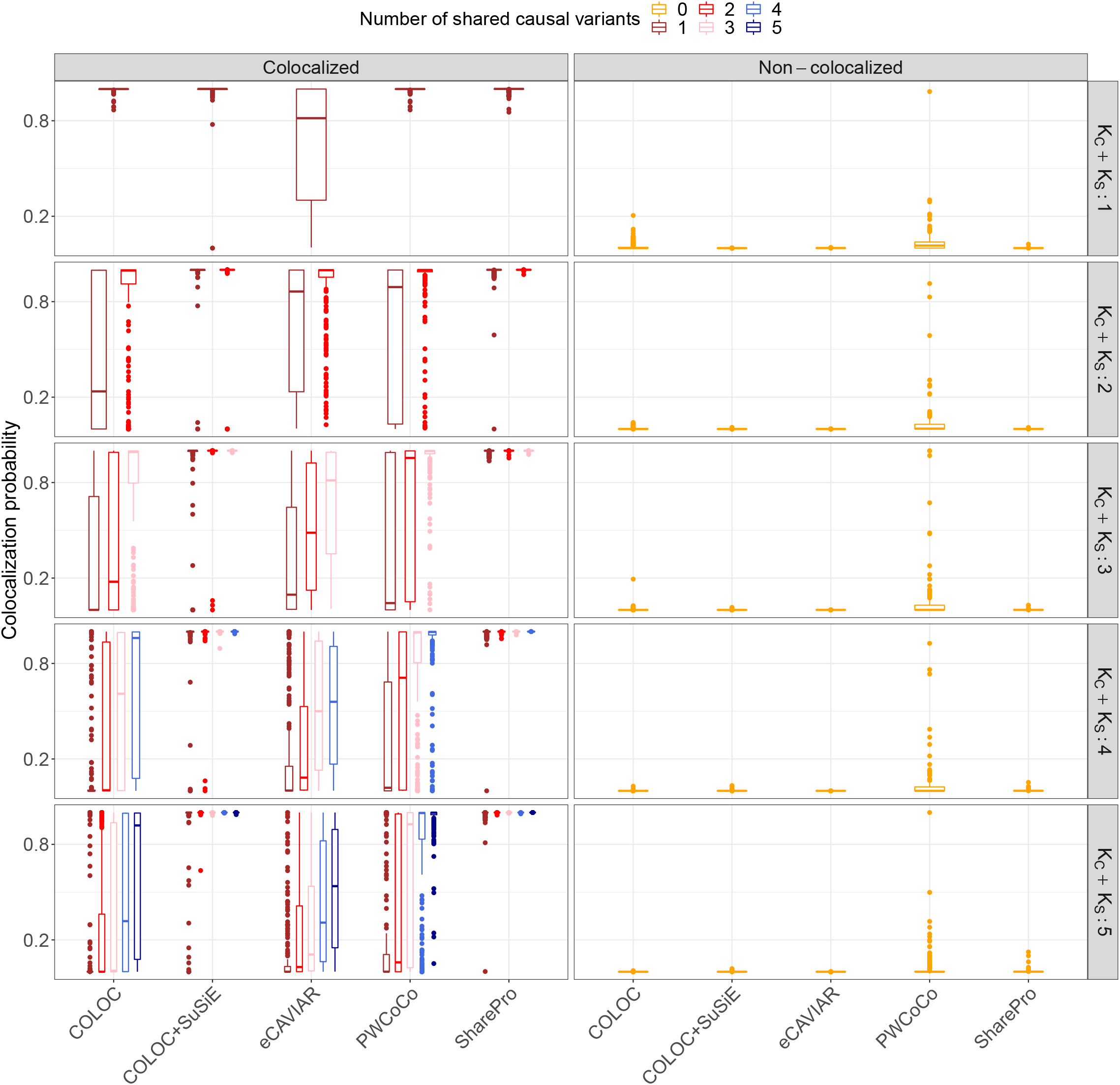
SharePro demonstrated improved power with a well controlled false positive rate for colocalization analysis. Colocalization probabilities derived by five methods based on 50 replicates in each of the five loci are illustrated. Rows represent the different numbers of causal variants (*K*_*C*_ + *K*_*S*_) and colors represent the different numbers of shared causal variants (*K*_*C*_) between the two simulated traits. Median colocalization probabilities across a total of 250 replicates are indicated by horizontal bars and inter-quartile ranges are represented by boxes.

**Figure 3.**
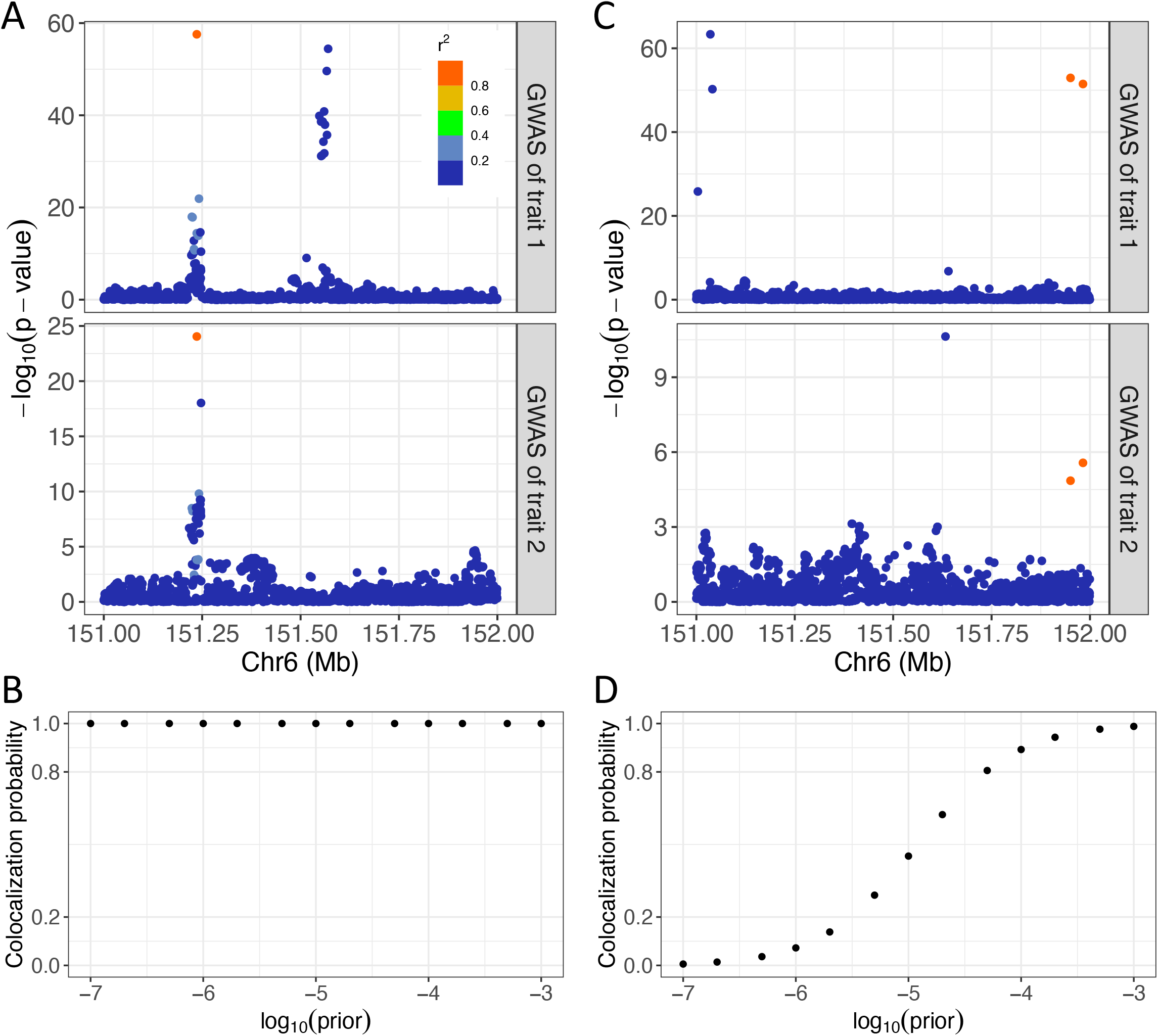
Prior sensitivity analysis in SharePro. (A) GWAS associations with a strong support for colocalization. Each dot represents a variant and the color indicates its correlation with the simulated colocalized variant. (B) Prior sensitivity analysis in the case of a strong support for colocalization. The x-axis stands for prior colocalization probabilities in the logarithmic scale and the y-axis stands for posterior colocalization probabilities. (C) GWAS associations with a weak support for colocalization. Each dot represents a variant and the color indicates its correlation with the simulated colocalized variant. (D) Prior sensitivity analysis in the case of a weak support for colocalization. The x-axis stands for prior colocalization probabilities in the logarithmic scale and the y-axis stands for posterior colocalization probabilities.

### 2.2 SharePro for colocalization analysis

In SharePro, we assume there are altogether *K* effect groups (for either trait **y**_1_ or trait **y**_2_, or both) in a locus with *G* variants. Similar to our previous work on the sparse projection formulation of the SuSiE model [15, 18, 19], for the *k*^*th*^ (*k* ∈ {1, …, *K*}) effect group, SharePro uses **s**_*k*_, a sparse indicator shared by both traits to specify its variant representations (**Figure 1**). This indicator follows a multinomial distribution:

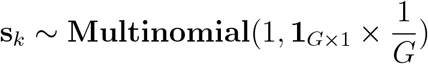

We use two additional sets of trait-specific variables to describe relationships between the *k*^*th*^ effect group and each trait: causal indicators *c*_*k*1_, *c*_*k*2_ of whether the *k*^*th*^ effect group is causal for **y**_1_ and **y**_2_ and *β*_*k*1_ and *β*_*k*2_ for their corresponding effect sizes (here we illustrate the model with two traits but it is also compatible with multiple traits):

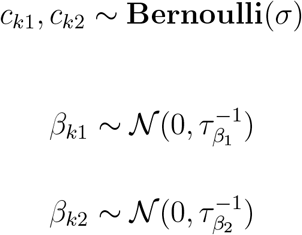

Denoting the genotype matrix as **X**_1_ and **X**_2_, for traits **y**_1_ and **y**_2_, we have:

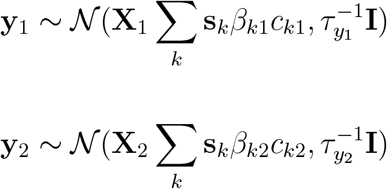

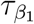 and 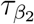 are hyperparameters for effect sizes while τ_*y*1_ and τ_*y*2_ are hyperparameters for residual variances; *σ* is the important hyperparameter for prior colocalization probability. We discuss choices of these hyperparameters in the **Supplementary Notes**. The colocalization probability for the *k*^*th*^ effect group is represented by the posterior probability of *p*(*c*_*k*1_ = *c*_*k*2_ = 1|**y**_1_, **y**_2_, **X**_1_, **X**_2_). We use an efficient variational inference algorithm [18, 20, 21] adapted for GWAS summary statistics for posterior inference (detailed in the **Supplementary Notes**).

### 2.3 Simulation studies

We conducted simulation studies under different causal settings to evaluate the performance of colocalization methods. We randomly sampled five 1-Mb loci from the genome and extracted their genotypes for 25,000 and 1,000 non-overlapping UK Biobank European ancestry individuals [22] to simulate trait 1 and trait 2, respectively. For each locus, we calculated the LD matrix using PLINK [23].

In each locus, we randomly sampled *K*_*C*_ causal variants to be shared across traits and additionally *K*_*S*_ causal variants to be specific for each trait. For example, when *K*_*C*_ = 0 and *K*_*S*_ = 1, there was one causal variant for trait 1 and one different causal variant for trait 2; When *K*_*C*_ = 1 and *K*_*S*_ = 0, there was one causal variant shared by both traits. We set the per-variant heritability to be 0.01 in trait 1 and 0.05 in trait 2. With simulated traits, we performed GWAS using GCTA [24] to obtain summary statistics. We repeated this process 50 times for each setting.

With LD information and simulated summary statistics, we performed colocalization analysis with five different methods (**Table 1**) using a default prior colocalization probability of 1 *×* 10^−5^ and obtained posterior colocalization probabilities from COLOC [1]. Both COLOC+SuSiE [4] and PWCoCo [16] generated multiple pairs of colocalization probabilities, with the maximum used as colocalization probabilities. For eCAVIAR, we also used the maximum variant-level colocalization probabilities as locus-level colocalization summary [2]. Similarly in SharePro, maximum colocalization probabilities across all identified effect groups were used.

**Table 1:** Summary of colocalization method features.

| Method | Multiple causal variants | Signal identification | Posterior summary | Running time (second; SD) | Reference |
| --- | --- | --- | --- | --- | --- |
| COLOC | X | X | locus-level | 0.1 (0.1) | [1] |
| COLOC+SuSiE | ✓ | separate fine-mapping | paired locus-level | 14.0 (3.3) | [5] |
| eCAVIAR | ✓ | separate fine-mapping | variant-level | 227.7 (89.3) | [2] |
| PWCoCo | ✓ | conditional analysis | paired locus-level | 38.1 (20.5) | [8] |
| SharePro | ✓ | joint fine-mapping | effect group-level | 4.3 (1.1) | this study |

A colocalization probability *>* 0.8 was considered strong evidence supporting colocalization, while a colocalization probability *<* 0.2 was considered evidence against colocalization [3].

### 2.4 Colocalization analysis of RSPO3 pQTL and eBMD GWAS

We examined the utility of SharePro by assessing the colocalization between a cis-pQTL locus of the circulating abundance of RSPO3, and a GWAS locus identified for eBMD using heel quantitative ultrasound measurement. We obtained UK Biobank eBMD GWAS summary statistics from the GEFOS consortium [25] and RSPO3 pQTL summary statistics from the Fenland study [26]. The LD matrix was calculated using UK Biobank European ancestry individuals and colocalization analysis was performed with five different methods (**Table 1**) using a default prior colocalization probability of 1 *×* 10^−5^.

## 3 Results

### 3.1 Simulation studies

To evaluate the performance of SharePro in colocalization analysis, we performed simulations under different causal settings. SharePro achieved the highest power in most settings. Specifically, in the simple scenario of only one causal variant (*K*_*C*_ + *K*_*S*_ = 1), COLOC, PWCoCo and SharePro accurately identified all simulated cases of colocalization with a colocalization probability above 0.8 (**Figure 2** and **Supplementary Table** S1). Meanwhile, COLOC + SuSiE only identified 98.8% cases of colocalization (**Supplementary Table** S1) while the locus-level colocalization summary derived from eCAVIAR only identified 51.2% of the simulated cases of colocalization (**Supplementary Table** S1).

In more challenging scenarios with multiple causal variants, SharePro maintained the highest power for colocalization analysis, followed by COLOC + SuSiE. For example, with *K*_*C*_ = 1 and *K*_*S*_ = 1 and a colocalization probability cutoff at 0.8, SharePro achieved a true positive rate of 99.2%, while the second best method COLOC + SuSiE achieved a true positive rate of 97.6% (**Figure 2 and Supplementary Table** S1). In contrast, as expected, since the one-causal-variant assumption was not satisfied, the performance of COLOC became worse and only identified 44.4% cases of colocalization (**Supplementary Table** S1). With more than one causal variant shared between the two simulated traits (*K*_*C*_ *>* 1), SharePro consistently identified all cases of colocalization and outperformed other methods (**Figure 2** and **Supplementary Table** S1).

When causal variants were different across the simulated traits (non-colocalized), the colocalization probabilities obtained by COLOC, COLOC+SuSiE, eCAVIAR and SharePro were consistently below 0.2 (**Figure 2** and **Supplementary Table** S2). In contrast, PWCoCo had higher colocalization probabilities. For instance, with *K*_*C*_ = 0 and *K*_*S*_ = 1, PWCoCo had a false positive rate of 2.4% with a colocalization probability cutoff at 0.2 (**Supplementary Table** S2).

Moreover, SharePro also demonstrated high computational efficiency (**Table 1)**. Across different simulation settings, on average, SharePro took 4.3 seconds to assess colocalization in a 1-Mb locus, which was only longer than COLOC. In contrast, on average, eCAVIAR took more than 3 minutes to assess colocalization in the same locus **(Table 1)**.

We additionally performed prior sensitivity analysis **(Supplementary Notes**) to examine the impact of prior colocalization probabilities on posterior colocalization probabilities and showcased two representative scenarios in **Figure 3**. When the GWAS summary statistics demonstrate strong colocalization pattern **(Figure 3A**), varying prior colocalization probabilities does not drastically change the posterior colocalization probabilities **(Figure 3B**). In contrast, when statistical evidence from GWAS associations is weak **(Figure 3C**), the posterior colocalization probabilities increases with the prior colocalization probabilities **(Figure 3D**).

### 3.2 RSPO3-eBMD example

The eBMD measured at the heel using quantitative ultrasound is an important biomarker of osteoporosis and strongly predicts fracture risk [25, 27, 28]. RSPO3 is a known modulator of the Wnt signaling pathway that plays a crucial role in maintaining bone homeostasis [29, 30]. It has been experimentally shown that the abundance of RSPO3 strongly influences the proliferation and differentiation of osteoblasts and regulates bone mass [13]. Therefore, it is biologically plausible that the cis-pQTL of RSPO3 colocalize with an eBMD GWAS locus.

However, although the marginal genetic associations for RSPO3 abundance and eBMD demonstrated a highly similar pattern (**Figure 4A)**, existing methods indicated no or minor evidence of colocalization **(Figure 4B)**. With SharePro, we identified multiple effect groups in this region and colocalization results indicated that both rs7741021/rs9482773 and rs853974 were shared causal signals between circulating RSPO3 abundance and eBMD **(Supplementary Table** S3). We explored different hyperparameter settings for prior colocalization probabilities in SharePro (**Supplementary Notes**) and obtained robust colocalization results (**Supplementary Tables** S4-7).

## 4 Discussion

In this work, we present SharePro to integrate LD modelling and colocalization assessment that extends the classical COLOC framework to account for multiple causal signals. Compared to methods that adopt a two-step strategy to relax the one-causal-variant assumption in COLOC, the effect group-level approach in SharePro can effectively reduce the impact of LD in aligning causal signals, resulting in improved power for colocalization analysis. Under different simulation settings, SharePro achieved the highest power with a well-controlled false positive rate. Additionally, SharePro also demonstrated high computational efficiency.

Through the example of the colocalization analysis of RSPO3 cis-pQTL and eBMD GWAS, we demonstrated that SharePro could correctly identify biologically plausible colocalization in the presence of multiple causal signals. In the RSPO3 locus, both the RSPO3 pQTL study and the eBMD GWAS are well-powered and the marginal associations exhibit a similar pattern (**Figure 4B**). However, the lead variants with the smallest marginal p-value in this locus, although highly correlated, are different for circulating RSPO3 abundance and eBMD (Figure 4B). In the presence of multiple causal signals, colo-calization analysis in this locus using existing methods has been challenging. In SharePro, correlated variants are grouped into effect groups and as a result, the impact of misaligned lead variants on colocal-ization analysis is mitigated.

**Figure 4.**
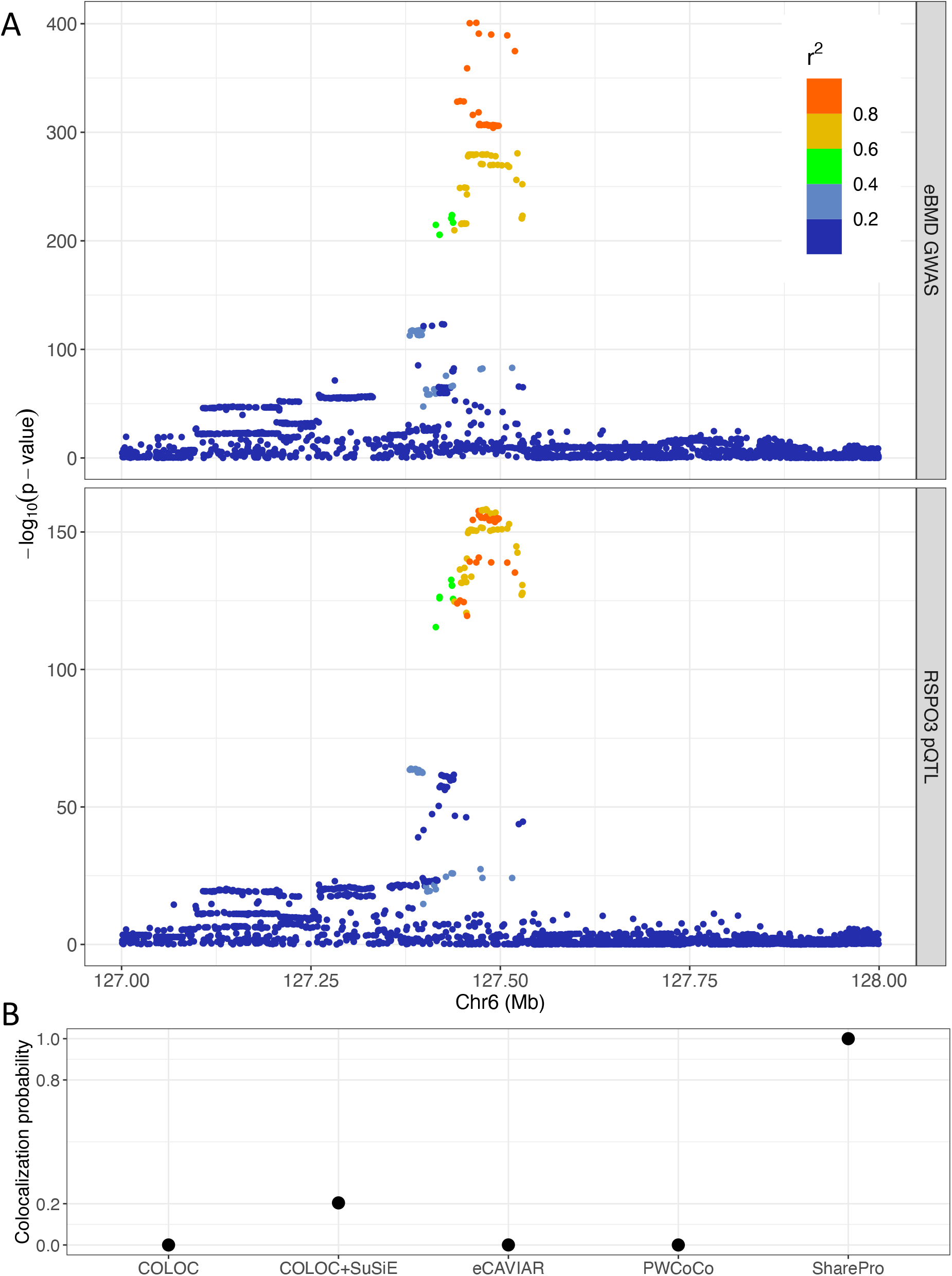
SharePro identified shared effect groups between RSPO3 pQTL and eBMD GWAS. (A) Marginal associations from eBMD GWAS and RSPO3 pQTL. The x-axis indicates chromosome and position information and the y-axis represents p-value on the logarithmic scale. Each dot represents a variant and its color indicates its correlation (*r*^2^) with the colocalized variant rs7741021. (B) Colocalization probabilities assessed by different colocalization methods for RSPO3 pQTL and eBMD GWAS.

An important hyperparameter in colocalization analysis is the prior colocalization probability. In SharePro, the default prior colocalization probability is 1 × 10^−5^. In COLOC, this hyperparameter is represented as *p*_12_ with a default value of 1 × 10^−5^ [1]. Because the prior colocalization probability can impact posterior colocalization probability, especially when GWAS are not well-powered, it is necessary to explore a range of different values to evaluate the robustness of colocalization results [4].

There are other cautions in colocalization analysis that also apply to SharePro. First, summary statistics-based analysis requires that the LD reference panel matches the LD structure underlying the samples included in GWAS. In SharePro, LD mismatch can lead to convergence issues for the algorithm. Second, the validity of colocalization results relies on the rigor of GWAS in carefully accounting for population stratification and other unmeasured confounding factors. Variants associated with shared confounding factors can also be considered colocalized. Third, the power to detect colocalization is dependent on the power of fine-mapping. We strongly suggest that prior sensitivity analysis should be performed to evaluate whether the GWAS are well-powered for colocalization analysis.

In summary, we have developed SparsePro to extend COLOC for colocalization analysis. With increased power and well-controlled false positive rate at a low computation cost, SharePro is suitable for large-scale colocalization analysis. With the increasing number of publicly available GWAS summary statistics, we envision SharePro will have the potential to substantially deepen our understanding of complex traits and diseases.

## 6 Supporting Information

### 6.1 Supplementary Notes

### 6.2 Supplementary Tables

## 7 Data and Software Availability

The SharePro software for colocalization analysis is openly available at https://github.com/zhwm/SharePro_coloc and the analysis conducted in this study is available at https://github.com/zhwm/SharePro_coloc_analysis. GWAS summary statistics for eBMD was obtained from the GEFOS consortium at http://www.gefos.org. GWAS summary statistics for RSPO3 pQTL was obtained from the Fenland study at https://omicscience.org/apps/pgwas/. Both COLOC and COLOC+SuSiE are included in the coloc (version 5.1.0) R package obtained from CRAN. PWCoCo was obtained from GitHub at https://github.com/jwr-git/pwcoco. eCAVIAR was obtained from GitHub at https://github.com/fhormoz/caviar.

## 8. Acknowledgements

W.Z. has been supported by a doctoral training fellowship from the FRQNT (319188) and the Healthy Brains, Healthy Lives Program, funded by the Canada First Research Excellence Fund (CFREF), Quebec’s Ministère de l’É conomie et de l’Innovation (MEI), and the Fonds de recherche du Québec (FRQS, FRQSC and FRQNT). T.L. has been supported by a Schmidt AI in Science Postdoctoral Fellowship and a Vanier Canada Graduate Scholarship. Y.L. is supported by Natural Sciences and Engineering Research Council (NSERC) Discovery Grant (RGPIN-2019-0621), Fonds de recherche Nature et technologies (FRQNT) New Career (NC-268592), and Canada First Research Excellence Fund Healthy Brains for Healthy Life (HBHL) initiative New Investigator start-up award (G249591). H.S.N holds a Canada Research Chair funded by the Canadian Institutes of Health Research and has been supported by NSERC Discovery Grant (RGPIN-2018-05962). This research used the NeuroHub infrastructure and was undertaken thanks in part to funding from the Canada First Research Excellence Fund, awarded through the Healthy Brains, Healthy Lives initiative at McGill University. This research was enabled in part by support provided by Calcul Québec and the Digital Research Alliance of Canada. This research has been conducted using the UK Biobank Resource under Application Number 45551.

## 9 Author contributions

Conceptualization: W.Z.; Data curation: W.Z., T.L.; Formal analysis: W.Z.; Funding acquisition: W.Z., Y.L., H.S.N, J.D.; Investigation: W.Z.; Methodology: W.Z.; Project Administration: R.S., H.S.N and J.D.; Resources: Y.L., H.S.N and J.D.; Software: W.Z.; Supervision: Y.L., R.S., H.S.N and J.D.; Validation: W.Z., T.L., R.S., Y.L., H.S.N and J.D.; Visualization: W.Z.; Writing – Original Draft Preparation: W.Z.; Writing – Review & Editing: W.Z., T.L., R.S., Y.L., H.S.N and J.D.

## 10 Disclosures

T.L. is an employee of 5 Prime Sciences Inc. The other authors have no relevant disclosures.

## SharePro Supplementary Notes

### 1 A variational inference algorithm for Bayesian colocalization

In SharePro (**Figure 1**), similar to our previous work on the sparse projection formulation of the SuSiE model [15,18,19], with a shared projection matrix **S**_*G×K*_ = [**s**_1_, …, **s**_*K*_], we can group correlated variants into *K* effect groups where

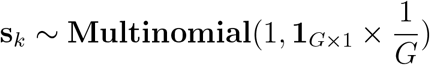

is the sparse indicator for the variant compositions in the *k*^*th*^ effect group. We have trait-specific indicator vectors **c**_1_ = [*c*_11_, …, *c*_*K*1_] and **c**_2_ = [*c*_12_, …, *c*_*K*2_] to characterize the causal statuses of effect groups on traits where

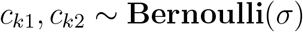

With trait-specific effect size vectors **β**_1_ = [*β*_11_, …, *β*_*K*1_] and **β**_2_[*β*_12_, …, *β*_*K*2_] where

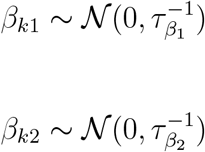

and denoting the genotype matrix as **X**_1_ and **X**_2_, for traits **y**_1_ and **y**_2_, we have:

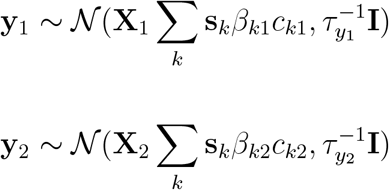

In colocalization analysis, we are interested in the posterior probabilities of causal indicators based on the observed traits **y**_1_, **y**_2_ and the genotypes **X**_1_ and **X**_2_. Inference of the exact posterior distribution of causal indicators **c**_1_, **c**_2_ and variant representations in effect groups **S** is difficult. Similar with the IBSS algorithm [15] proposed by SuSiE and our previous work on paired mean field variational inference algorithm [18, 21], we use a paired mean field factorized variational family [21]

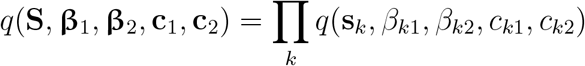

to approximate the desired posterior distribution:

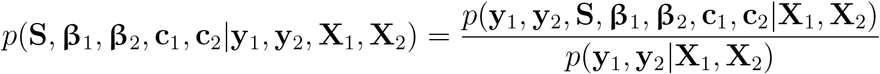

We can obtain the optimal approximation by maximizing the evidence lower bound (ELBO) [20]:

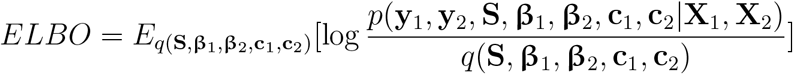

with the following conditions satisfied for each *k* [20]:

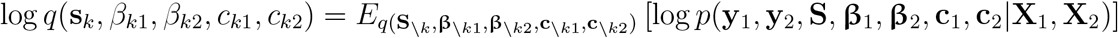

*where* 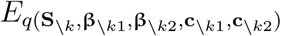 is the expectation with respect to the variational distribution excluding the *k*^*th*^ component. If we write out the joint probability:

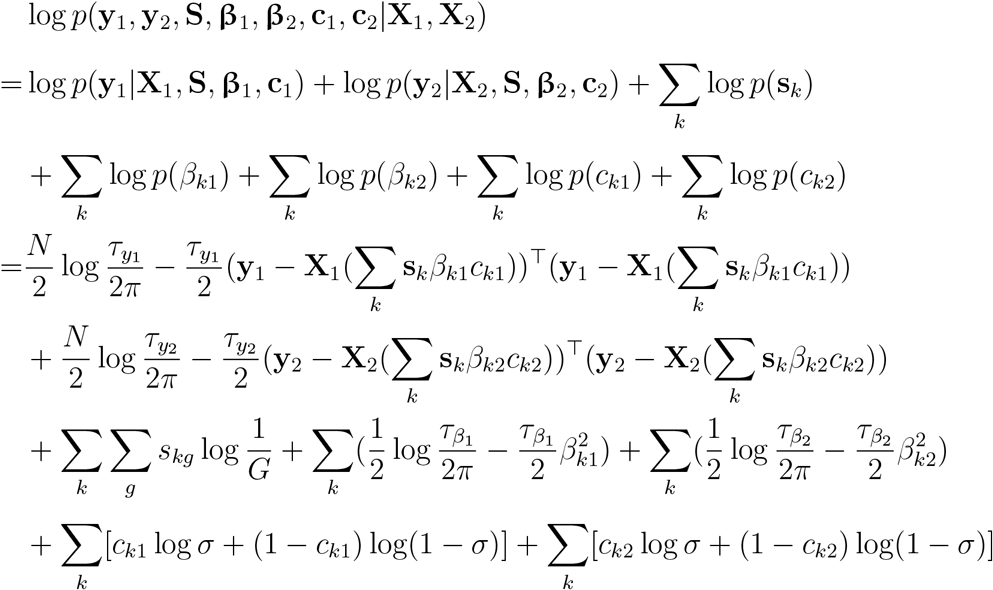

and denote 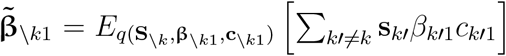 and 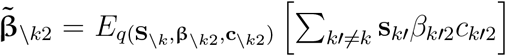 the required conditions can be simplified into four different cases for the *k*^*th*^ effect group:

**Case 1**: *c*_*k*1_ = 0 = *c*_*k*2_ = 0, i.e. the *k*^*th*^ effect group is not causal for either trait:

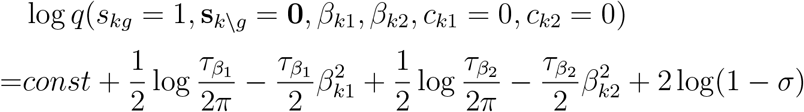

After integrating out *β*_*k*1_ and *β*_*k*2_, we have:

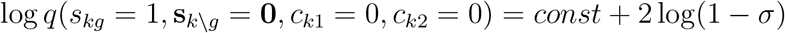

**Case 2 (trait 1 specific)**: *c*_*k*1_ = 1 and *c*_*k*2_ = 0, i.e. the *k*^*th*^ effect group is causal for trait **y**_1_ but not for trait **y**_2_:

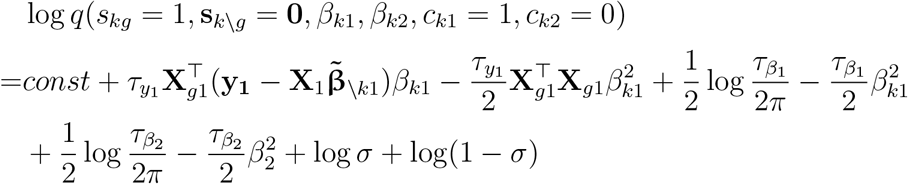

We recognize that 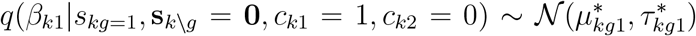. By matching sufficient statistics of the normal distribution, we have variational parameters for *β*_*k*1_:

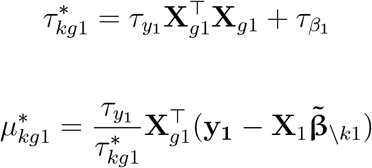

After integrating out *β*_*k*1_ and *β*_*k*2_, we have:

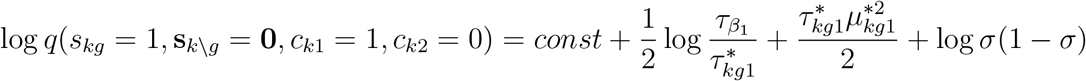

**Case 3 (trait 2 specific)**: *c*_*k*1_ = 0 and *c*_*k*2_ = 1, i.e. the *k*^*th*^ effect group is causal for trait **y**_2_ but not for trait **y**_1_:

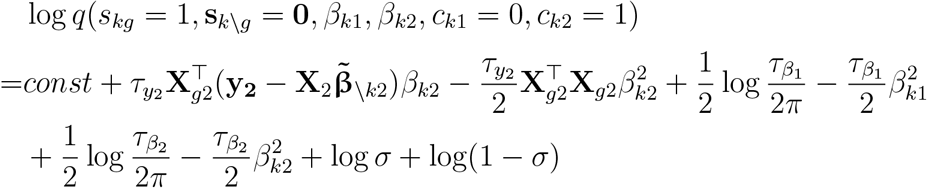

Similarly, we recognize that 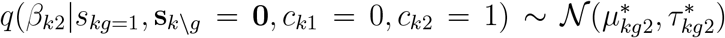. By matching sufficient statistics for the normal distribution, we can obtain the following variational parameters for *β*_*k*2_:

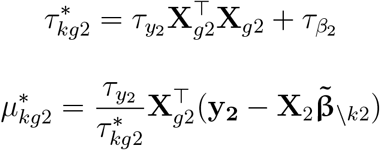

After integrating out *β*_*k*1_ and *β*_*k*2_, we have:

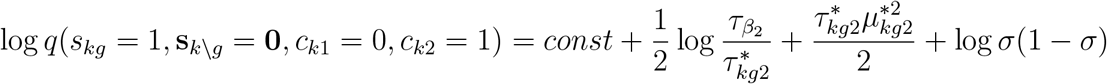

**Case 4 (colocalization)**: *c*_*k*1_ = 1 = *c*_*k*2_ = 1, i.e. the *k*^*th*^ effect group is causal for both trait **y**_1_ and trait **y**_2_:

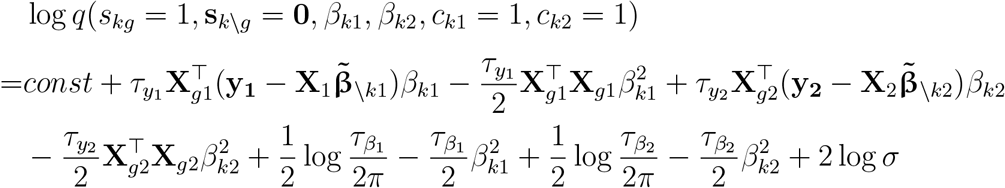

We recognize that 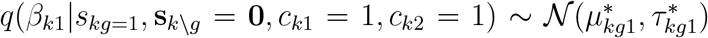 and 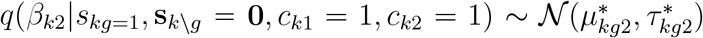. By matching sufficient statistics for these normal distribution, we obtain the following variational parameters for *β*_*k*1_ and *β*_*k*2_:

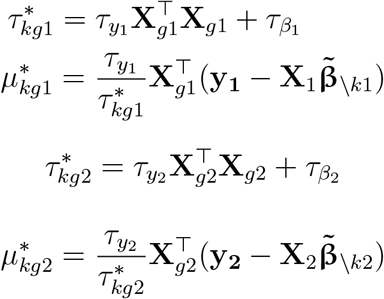

After integrating out *β*_*k*1_ and *β*_*k*2_, we have:

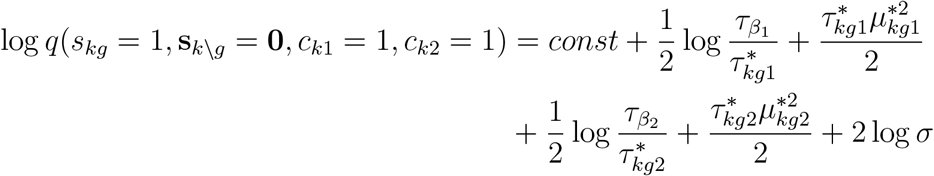

Combining all four cases, we have the conditional distributions for *c*_*k*1_ and *c*_*k*2_:

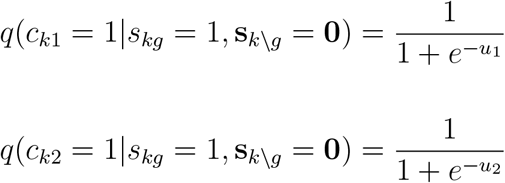

Where

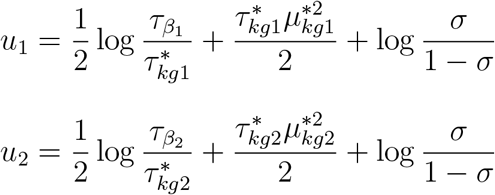

After integrating out *c*_*k*1_ and *c*_*k*2_, we have the variational distribution for **s**_*k*_:

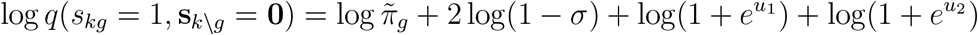

Therefore, for the *k*^*th*^ effect groups, we can calculate the posterior colocalization probability as

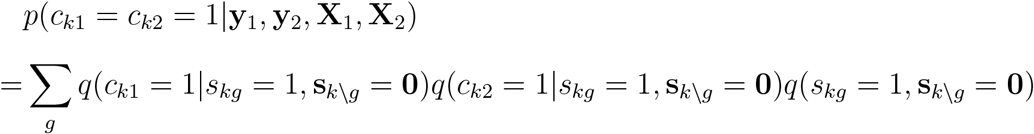

In summary, we have now derived **Algorithm** 1 **for colocalization analysis with SharePro:**

#### Algorithm 1: SharePro for genetic colocalization analysis

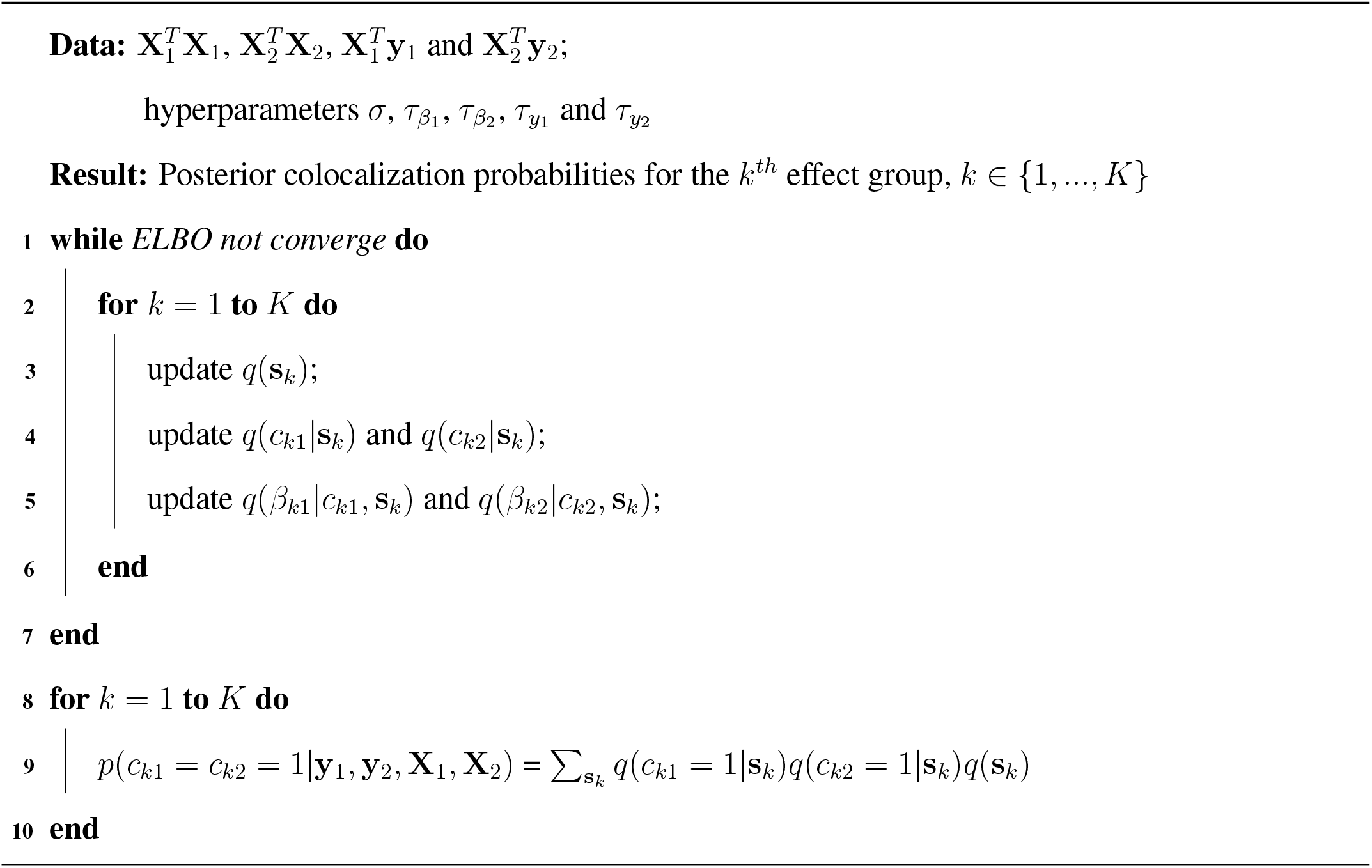

## 2 Adaptation to summary statistics

The information in individual-level data 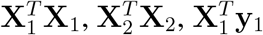 and 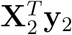 throughout the proposed **Algorithm** 1, which can be derived from GWAS summary statistics and a LD reference panel. Specifically, in most publicly available GWAS summary statistics, standardized effect sizes (z-scores) are usually available or can be derived from marginal effect sizes and standard errors. With standardized genotypes and phenotypes, we have:

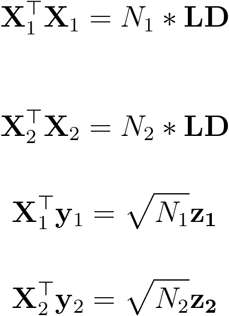

where *N*_1_ and *N*_2_ are sample sizes, **LD** is the variant Pearson correlation coefficient matrix and **z**_1_ and **z**_2_ are the z-scores in GWAS summary statistics for trait 1 and trait 2 respectively.

## 3 Hyperparameter estimation

Apart from the required quantities derived from GWAS summary statistics, there are also hyperparameters to be estimated in the colocalization algorithm: 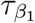 and 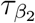 for effect size distributions, τ_y1_ and τ_y2_ for trait residual variances and *σ* for prior colocalization probability. As shown in our previous work [18], HESS-based heritability estimates [31] can provide suitable estimation for variance hyperparameters. Specifically, we can obtain the local heritability (*ĥ*^2^) in a locus as well as per-variant heritability 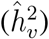 with the HESS [31] estimator using GWAS summary statistics, and use them to set hyperparameters: 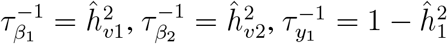 and 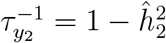.

An important hyperparameter in Bayesian colocalization is the prior colocalization probability *σ*. We set its default value to 1 *×* 10^−5^ (the same default value as used in COLOC). However, the impact of prior colocalization probabilities on posterior colocalization probabilities depends on the power of GWAS. In simulation studies, we explored a range of prior: 1 *×* 10^−7^, 2 *×* 10^−7^, 5 *×* 10^−7^, 1 *×* 10^−6^, 2 *×* 10^−6^, 5 *×* 10^−6^, 1 *×* 10^−5^, 2 *×* 10^−5^, 5 *×* 10^−5^, 1 *×* 10^−4^, 2 *×* 10^−4^, 5 *×* 10^−4^, 1 *×* 10^−3^ and showcased two representative simulation examples in **Figure 3**.

## Notes

https://github.com/zhwm/SharePro_coloc

https://github.com/zhwm/SharePro_coloc_analysis

